# *Alternaria solani* infection reprograms potato leaf metabolism and highlights potential defence and metabolic markers

**DOI:** 10.64898/2026.08.17.745268

**Authors:** Portia D Singh, Ramnath Nayak, Sanjeev Sharma, Shyam Kumar Masakapalli

**Affiliations:** School of Biosciences and Bioengineering, Indian Institute of Technology Mandi, Kamand, Himachal Pradesh-175075, India; ICAR-Central Potato Research Institute, Shimla, Himachal Pradesh, India

**Keywords:** Potato (*Solanum tuberosum* L.), Early blight, *Alternaria solani*, Plant-microbe interaction, Metabolic pathways, Nitrogen remobilisation Metabolite biomarkers, Lesion development, Disease diagnostics, GC-MS, Metabolomics

## Abstract

Potato (*Solanum tuberosum* L.), the world’s fourth most cultivated crop, suffers yield losses of up to 40-50% from early blight caused by the necrotrophic fungal pathogen *Alternaria solani*. In this study we performed gas chromatography-mass spectrometry (GC-MS)-based untargeted metabolomics to characterize temporal alterations in metabolite composition, metabolic pathway regulation, and discriminatory biomarker metabolites in the susceptible Indian potato variety *Kufri Jyoti*, analyzing infected leaves, non-infected leaves, and lesion-associated necrotic tissues across four days post-inoculation (DPI).Metabolite annotation identified 58 compounds, including sugars, organic acids, amino acids, and secondary metabolites.. Multivariate analyses resolved distinct, largely non-overlapping metabolic clusters for control, infected leaves (1-4 DPI), and lesion tissue (Bs1-Bs3). A biphasic metabolic response was observed: early infection (1-2 DPI) was characterized by general suppression of primary metabolism, while late infection (3-4 DPI) showed pronounced upregulation of glycolysis, the TCA cycle, GS/GOGAT, and the shikimate pathway. Key discriminatory metabolites included asparagine, oxoproline, GABA, phenylalanine, and aromatic amino acids. Lesion tissues exhibited distinct metabolic fingerprints, with early disruption of amino acid recycling followed by a late rebound of defense-associated metabolites. Notably, defence-associated phenolics were detected exclusively within lesion tissue and were absent from whole-leaf profiles, demonstrating that spatially resolved lesion sampling captures defence chemistry that whole-leaf analysis alone would miss. The identified biomarker metabolites, particularly those linked to the shikimate and GS/GOGAT pathways, represent promising candidates for metabolite-assisted breeding and targeted crop protection strategies against early blight in potato.

**Graphical Abstract:** 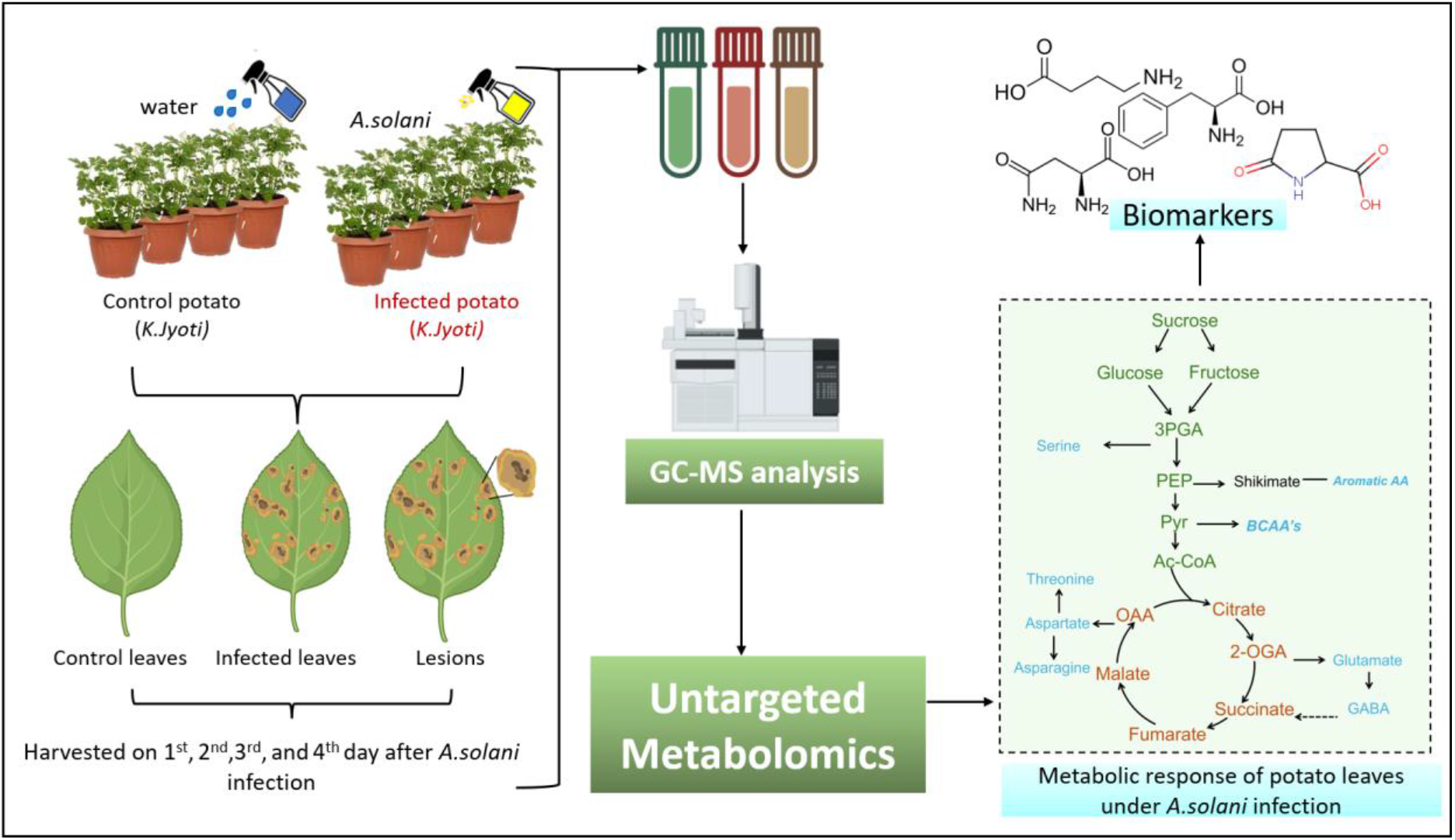

## Introduction

Potato (*Solanum tuberosum* L.) ranks as one of the world’s most economically significant tuberous crops and, due to rising global food demand, has become the fourth most widely cultivated crop worldwide, following wheat, rice, and maize (Singh et al., 2024). Rich in immunity-boosting biomolecules including vitamins C and B6, potassium, and phenolic compounds, potato is an indispensable contributor to global nutrition and food security (Zaheer & Akhtar, 2016). However, potato productivity is threatened by both biotic and abiotic stresses that routinely cause significant yield losses (Kumar et al., 2023; Zhang, H.; et al., 2017a). Among these, early blight disease caused by the necrotrophic fungal pathogen *Alternaria solani* is one of the most destructive foliar diseases affecting potato and tomato worldwide, capable of causing yield reductions of 40-50% under conducive field conditions (Adhikari et al., 2017; Attia et al., 2020; Sharma et al., 2025). Warm, humid climates and nitrogen-deficient soils are particularly favorable to disease outbreaks, with severe losses documented across major potato-growing regions including South Asia (Niu et al., 2022).

Management of early blight has relied on the development of resistant cultivars alongside the application of chemical fungicides (Sharma et al., 2017). While fungicides remain the most effective control strategy, their prolonged and repeated use has driven the emergence of *A. solani* strains with resistance to multiple fungicide classes including quinone outside inhibitors (QoIs), succinate dehydrogenase inhibitors (SDHIs), and anilinopyrimidines in field populations across several countries (Budde-Rodriguez et al., 2022; Bauske & Gudmestad, 2018). This increasing resistance problem, combined with environmental concerns over agrochemical accumulation in soil and water, highlights the urgent need to develop sustainable, knowledge-based disease management strategies for early blight control (Li et al., 2023).

Plant immunity against necrotrophic pathogens involves a complex network of metabolic and molecular responses. Following pathogen invasion, plants rapidly perceive the invading organism and activate signalling pathways that coordinate defence responses (Peyraud et al., 2017). As infection progresses, these signalling events reprogram both primary and secondary metabolism, leading to the accumulation of metabolites that either inhibit pathogen growth or strengthen plant defence (Piasecka et al., 2015). Secondary metabolites, including phenylpropanoids, terpenoids, and oxylipins, play central roles in pathogen resistance, plant fitness, and adaptation to environmental stress (Yang et al., 2018). Therefore, identifying the metabolic pathways that are activated or suppressed during infection is essential for understanding disease resistance and for discovering biomarkers that could support improved crop protection strategies (Botero et al., 2018). Metabolomics has become an important tool for investigating these metabolic changes because it provides a comprehensive view of the biochemical responses occurring during plant-pathogen interactions. When combined with multivariate statistical analysis and bioinformatics, metabolomics can identify metabolic pathways associated with disease progression and host defence (Serag et al., 2023). Studies of necrotrophic pathosystems have revealed common patterns of metabolic reprogramming across different host species. For example, *Botrytis cinerea* infection in *Arabidopsis* causes an early reduction in photosynthetic metabolism followed by activation of jasmonate and ethylene-mediated defence pathways (Windram et al., 2012). Likewise, *Alternaria alternata* infection in tomato induces phenylpropanoid biosynthesis and the accumulation of metabolites involved in scavenging reactive oxygen species (Singh et al., 2023a). Although these studies have provided valuable insights, defence responses remain highly dependent on the host species, cultivar, and pathogen, highlighting the need for crop-specific investigations. Among the available metabolomics platforms, gas chromatography-mass spectrometry (GC-MS) is widely used because it enables reliable detection of a broad range of primary metabolites, including amino acids, organic acids, sugars, sugar alcohols, and several secondary metabolites (Choudhury et al., 2021). GC-MS has been successfully applied to investigate potato responses to pathogens such as *Rhizoctonia solani* (Aliferis & Jabaji, 2012) and *Phytophthora infestans* (Zhu et al., 2022). However, despite the economic importance of early blight, a comprehensive kinetic GC-MS metabolomic analysis of potato leaves infected with *Alternaria solani* has not yet been reported. A few metabolomic studies have investigated *Alternaria* interactions using LC-MS-based platforms, primarily in tomato and the wild tomato species *Solanum cheesmaniae* (Singh et al., 2023a; Singh et al., 2023b). These studies provide valuable insights into host metabolic responses but do not address the temporal metabolic reprogramming of potato during *A. solani* infection. This represents an important knowledge gap, particularly because potato possesses a metabolic profile distinct from tomato, and disease progression by necrotrophic pathogens is likely to exhibit species-specific metabolic responses.

The present study addresses this gap by characterizing temporal changes in the metabolite composition of leaves from the susceptible Indian potato cultivar *Kufri Jyot*i following *A. solani* infection using GC-MS. Metabolic profiles were compared between healthy leaves and infected tissues (infection and necrotic lesion development) collected at different stages of disease progression. By identifying metabolites and metabolic pathways associated with disease progression, this study aims to improve our understanding of the potato-*A. solani* interaction and identify potential metabolic biomarkers of early blight. These findings may contribute to the development of more effective strategies for improving resistance to early blight in potato.

## 2. Materials and methods

### 2.1 Plant material and growth conditions

The potato variety (*Kufri Jyoti*) was cultivated in controlled conditions and inoculated with the pathogen. Potato seed tubers were provided by ICAR-Central Potato Research Institute, Shimla. Briefly, the seed tubers were sterilized by immersing them in a 2% sodium hypochlorite solution for 5 minutes. After drying, the seed tubers were planted in earthen pots measuring 20 × 20 × 14 cm. These pots contained a sterilized mixture of soil, cocopeat, perlite, and vermiculite in a ratio of 2:1:1:1 (by weight) and provided with optimal growing conditions within a controlled environment with the temperature adjusted to 22 °C,16 hrs of artificial light, and 70% relative humidity. Each experimental group comprised four biological replicates (individual plants), each representing an independent pot.

### 2.2 Pathogen inoculation

The fungal strain *Alternaria solani* (MTCC Accession No. 10690) was procured from the Microbial Type Culture Collection (MTCC), Chandigarh. The strain was cultured on potato dextrose agar (PDA) for 10 days to obtain sufficient sporulation. Spore suspensions were prepared by gently scraping the fungal culture using a sterile rubber spatula in the presence of Tween 20, followed by filtration through a double layer of sterile cheesecloth. Spore concentration was determined using a hemocytometer and adjusted to 4.6 × 10⁷ conidia/ml. Potato plants were inoculated at the early flowering stage (45 days post-planting) by spraying the spore suspension using a sterilized sprayer. Control plants were inoculated with double-distilled water (ddH₂O). After inoculation, plants were enclosed in polythene bags and incubated in the dark for 24 hours at >95% relative humidity (RH) to facilitate infection. Subsequently, the bags were removed, and RH was maintained at approximately 75% to promote disease progression. Leaf samples were collected at 1^st^, 2^nd^, 3^rd^, and 4^th^ days post inoculation (DPI), while lesion development (Bs) was monitored at 1^st^, 2^nd^, and 3^rd^ DPI for metabolomic analysis. At each sampling time point, four leaflets were harvested from four individual plants (n = 4). For lesion-stage samples (Bs1-Bs3), lesion tissues were similarly collected from four infected leaves. All samples were immediately flash-frozen in liquid nitrogen and stored at -80°C until further processing. Lesion tissues were not collected at the Bs4 stage because the leaves were completely colonized by *Alternaria solani*, making it impossible to distinguish lesion tissue from the surrounding infected leaf tissue.

### 2.3 Trypan blue staining

Trypan blue was employed as a vital stain to assess loss of cellular viability, following the method described by Koch et al., (1990). Potato leaves were collected at 1^st^ and 2^nd^ DPI and subjected to destaining in an acetic acid:ethanol solution (1:3, v/v) to remove chlorophyll. Subsequently, leaves were stained overnight in a lactophenol-trypan blue solution (0.01% Trypan Blue in a 1:1:1 [v/v/v] mixture of lactic acid, phenol, and double-distilled water). Excess stain was removed by rinsing the leaves with double-distilled water. Stained leaves were mounted on glass slides using 50% glycerol and examined microscopically for cell viability assessment.

### 2.4 Metabolites extraction and data acquisition using GC-MS

Lyophilized potato leaf tissue (∼50 mg) was used for soluble metabolite extraction following the protocol described by Lisec et al., (2006). Each sample was homogenized in 940 µL of extraction solvent comprising methanol:chloroform:water (3:1:1, v/v/v) and supplemented with 60 µL of ribitol (0.2 mg/mL in H₂O) as an internal standard. Samples were incubated at 70°C for 5 minutes in a thermomixer at 950 rpm, followed by centrifugation at 13,000 × g for 10 minutes at room temperature. Approximately 50 µL of the supernatant was transferred to a fresh microcentrifuge tube and vacuum dried. Dried extracts were derivatized via methoxyamine hydrochloride (MeOX) and trimethylsilyl (TMS) reagent treatment, as described by Masakapalli et al. (2013). Samples were subsequently analyzed using a GC-MS (GC ALS-MS 5977B, Agilent Technologies) equipped with HP-5ms (5% phenyl methyl siloxane) column (30 m × 250 μm × .25 μm) at IIT Mandi. Metabolite identification was based on comparison of mass ion fragments (m/z), retention times (RT), in house standards, and characteristic identifier ions with spectral data from the NIST 17 library (match factor ≥80%). For statistical and multivariate data analysis, we utilized MetaboAnalyst 6.0, accessible at https://www.metaboanalyst.ca/.. Data pre-processing included normalization by reference feature (ribitol internal standard), log₂ transformation, and pareto scaling. Differential abundance between conditions was assessed using one-way ANOVA with Tukey’s HSD post-hoc test; metabolites with a corrected p-value <0.05 and |log₂FC| ≥1.0 were considered significantly altered.

## 3. Results

### Development of early blight in potato leaves infected by *A. solani* infected

Disease progression following *Alternaria solani* inoculation of *Kufri Jyoti* leaves was tracked both externally, on whole leaves, and internally, via leaf cross-sections. Mock-inoculated control leaves remained symptomless throughout the observation period. Inoculated leaves showed the earliest visible external symptoms at the inoculation site by 1st day post-inoculation (DPI); lesions expanded visibly by 3^rd^ DPI, and by 4^th^ DPI leaves displayed extensive brown, necrotic lesions consistent with active colonisation (Figure 1.A). Microscopic imaging (Figure 1.B) showed that internal changes preceded this external progression. Control leaves showed no lesion formation. By 1^st^ DPI, early internal lesion development was already visible, ahead of the more pronounced external symptoms seen later. By 2^nd^ DPI, the internal lesion appeared fully established, indicating that substantial internal colonisation occurs within the first two days of infection, well before the external browning observed at 3-4 DPI (Figure 1.A). This time-course provided the rationale for sampling whole-leaf at 1-4 DPI and lesion tissue at sequential maturation stages (Bs1-Bs3) for the metabolite profiling described below. The viability of cells was assessed using the detached leaf assay and the Trypan blue test. The assay demonstrated that the potato leaves exhibited a higher rate of necrosis in leaf tissue after 2^nd^ day of infection, which was 2-fold higher than that of the 1^st^ day infection (Figure. 1.C).

**Figure 1:**
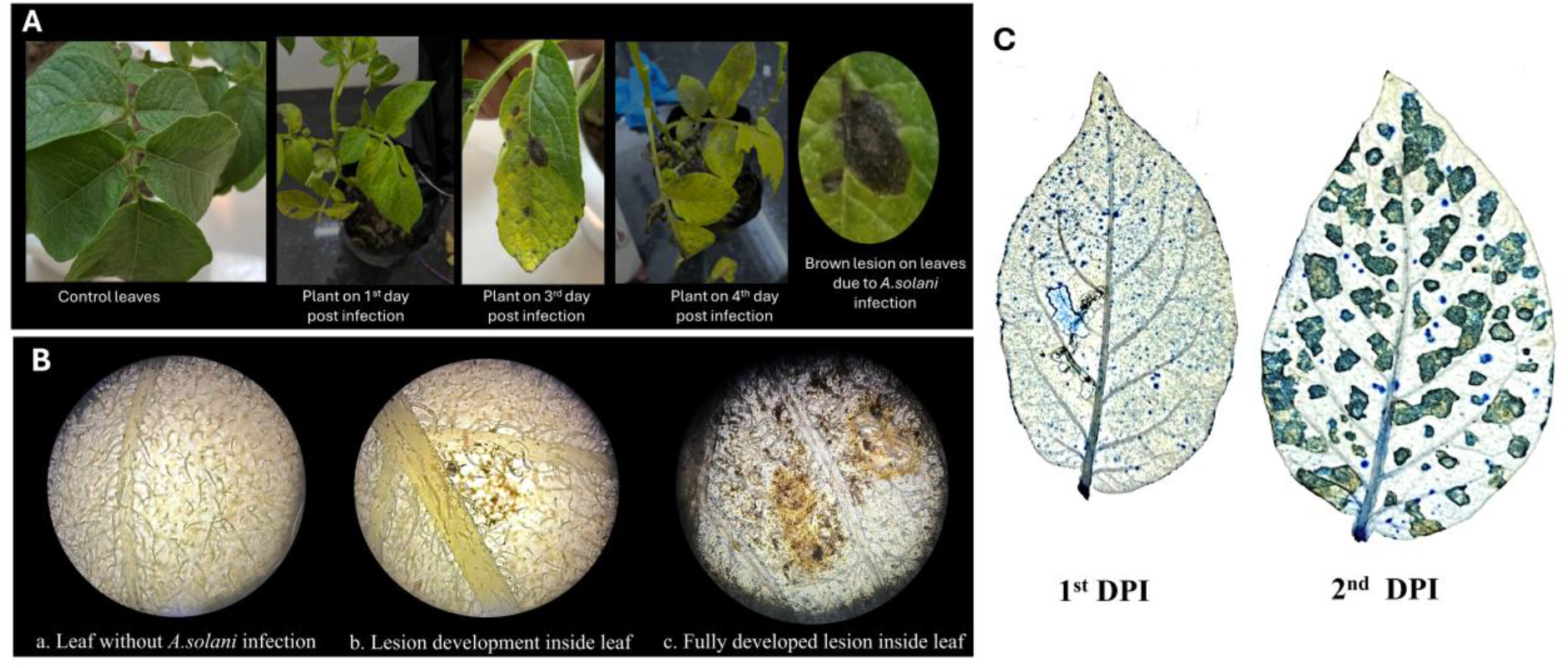
Morphological and microscopic changes captured in potato leaves (*Kufri Jyoti*) infected with *Alternaria solani* strain 10690, i.e., control and infected plants. Leaves were harvested at 1^st^,2^nd^,3^rd^, and 4^th^ days after inoculation. A) Morphological changes in potato leaves B) Microscopically captured potato leaf lesions demonstrating the development of *Alternaria solani* lesions within *Kufri Jyoti* plants infected with *Alternaria solani* C) Cell viability assay using Trypan blue staining in potato leaves of *K.Jyoti* variety 1^st^ and 2^nd^ day after post inoculation (DPI) with *Alternaria solani*.

### Comparative metabolomics elucidates early detection of *A. solani*-induced metabolic alterations in lesion versus whole-leaf sampling

Untargeted GC-MS profiling of the soluble extracts from *A. solani*-infected leaves was performed across two sampling schemes: the whole-leaf infection time course (1→4 DPI) and individual lesion developmental stages (Bs1→Bs3). 58 key metabolic features were identified and categorised as sugar alcohols, organic acids, amino acids, and secondary metabolites (Supplementary Table S1). The principle component analysis (PCA) of the whole-leaf tissue (control, 1-4 DPI) resolved five distinct, non-overlapping clusters (PC1 = 56.2%, PC2 = 32.9%), with control clearly separated from all infected time points along PC1 (Figure 2A). The early phase of infection (1-2 DPI) showed a drastic metabolic shift towards the positive PC2 axis, which later progressed towards the positive PC1 axis as the infection advanced (3-4 DPI). Further, the metabolites with the highest VIP scores (all >1.5) were asparagine, phenylalanine, oxoproline, GABA, and serine, followed by tyrosine, tryptophan, and galactinol (Figure 2C); all except GABA and dulcitol increased progressively from control through 3-4 DPI, while GABA and dulcitol were highest in control and declined with infection. A similar trend was observed in the PCA of the lesion tissues (Bs1-3), which resolved four groups (PC1 = 56.1%, PC2 = 32.7%; Figure 2B). Here, the Bs1 samples were visibly more dispersed than the other three groups, which formed tighter clusters, towards the negative PC1 axis. Across all eight conditions together, Bs2/Bs3 clustered apart from all other groups, Bs1 again showed the widest scatter, and 1-2 DPI and 3-4 DPI formed two adjoining but distinguishable clusters, with control positioned near the 1-2 DPI cluster (PC1 = 38.7%, PC2 = 24.9%; Supplementary Figure S1A). In the infection time-course, metabolites with the highest VIP scores (all >1.5) were asparagine, phenylalanine, oxoproline, GABA, and serine, followed by tyrosine, tryptophan, and galactinol (Figure 2C); all except GABA and dulcitol increased progressively from control through 3–4 DPI, while GABA and dulcitol were highest in control and declined with infection. In lesion-stage samples, gluconic acid, GABA, fucitol, glucose, and oxalic acid had the highest VIP scores (Figure 2D). Gluconic acid, GABA, glucose, and 1,5-anhydroglucitol were highest in control and lowest at Bs1, while fucitol, oxalic acid, galactaric acid, maltose, ribose, and sucrose were sharply and specifically elevated at Bs1 relative to control, Bs2, and Bs3. Consistent with these patterns dulcitol, fucitol, ribose, and galactaric acid as among the most discriminating metabolites with distinct accumulation patterns across infection and lesion stages (Supplementary Figure S1 B). Hierarchical clustering dendrogram analysis (Supplementary Figure S1 C) revealed that control, lesion day 1, and infection days 1 and 2 modulated their metabolomes similarly under *A. solani* infection, whereas infection on days 3 and 4 and lesions developed on days 2 and 3 modulated in a comparable pattern. In the infected leaves time-course heatmap (Supplementary Figure S1D), amino acids including valine, threonine, serine, asparagine, phenylalanine, tyrosine and isoleucine, along with galactinol and arbutin, were consistently elevated at 3-4 DPI relative to control and early time points. In the lesion-stage heatmap (Supplementary Figure S1E), sugars and sugar derivatives (ribose, maltose, mannobiose, fucitol), amino acids (tyrosine, threonine, phenylalanine, serine, oxoproline, valine), and phenolic/secondary metabolites (caffeic acid, chlorogenic acid, aucubin, salicylic acid) were elevated in Bs2 and Bs3 relative to CC and Bs1.

**Figure 2:**
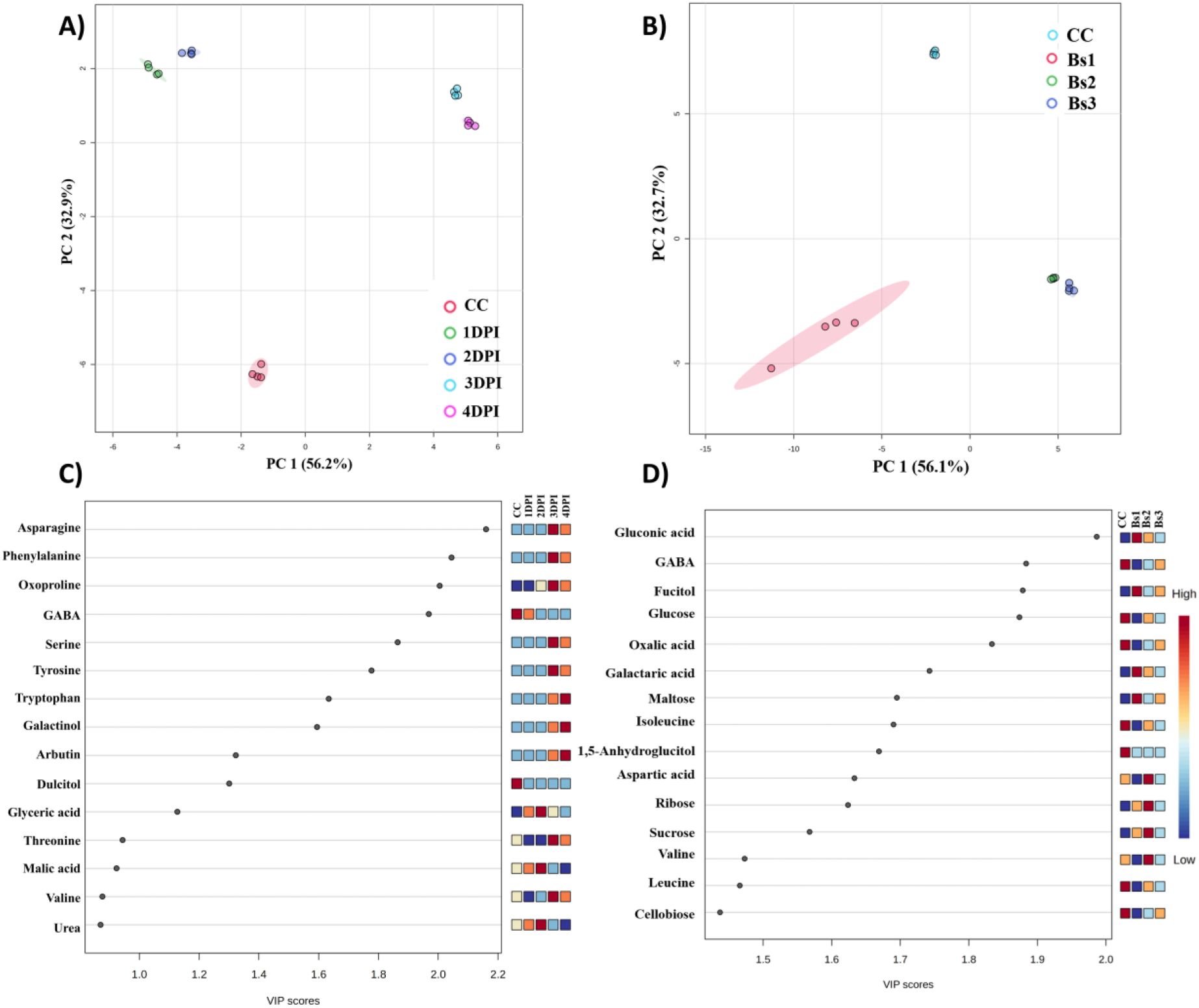
Metabolic variation captured in potato leaves infected with *A.solani* and in developed lesions through multivariate statistical analysis. A&B) PCA score plot the variation between control and infected leaves and control leaves and lesions developed. C&D) The VIP score plots display the significance of top 15 identified metabolites with a VIP score greater than 1. A trichromatic scale consisting of blue, yellow, and red colors is employed to visually represent the extent of a metabolite’s contribution to metabolic variation in different conditions. Abbreviations CC-controlled leaves; DPI: Days post infection; Bs: Lesion developed; numbers correspond to days post infection

### Metabolic reprogramming in potato leaves during *Alternaria solani* infection

Metabolite abundance is presented as log2 fold-change (Figure 3) relative to healthy controls, across a temporal axis (1-4 DPI, whole leaves) and a spatial axis (Bs1, lesion centre, through Bs3, lesion margin). Citrate and malate were depleted in both infected leaves and lesion tissue across nearly all time points/zones. Succinate and fumarate diverged: fumarate was elevated only at 3-4 DPI in leaves and unchanged in all lesion zones; succinate was elevated across all three lesion zones (increasing toward the lesion centre) but reached significance only transiently at 1 DPI in leaves (Supplementary Figure S2). GABA mirrored this divergence depleted throughout the leaf time-course, moderately depleted at Bs1-Bs2, and non-significant by Bs3.Valine, leucine, and isoleucine were depleted early (1-2 DPI) and accumulated at 3-4 DPI in leaves; in lesions, valine rose progressively from Bs1-Bs3 while leucine/isoleucine remained depleted or non-significant (Supplementary Figure S3). Aspartate and asparagine accumulated late in leaves (3-4 DPI); in lesions, aspartate was depleted at Bs1 before rising at Bs2-Bs3, while asparagine was elevated from Bs1 onward. Threonine and β-alanine behaved inversely to one another in leaves. Tyrosine, phenylalanine, and tryptophan showed the strongest, most consistent accumulation of any metabolite class elevated at 3-4 DPI in leaves and across all lesion zones, with tryptophan already elevated at Bs1. Chlorogenic acid, caffeic acid, and salicylic acid were detected only in lesion tissue, increasing progressively from Bs1 to Bs3.Serine accumulated from 3 DPI (leaves) and from Bs1 (lesions); glycine, measured only in the lesion, was elevated from Bs2 onward.

**Figure 3:**
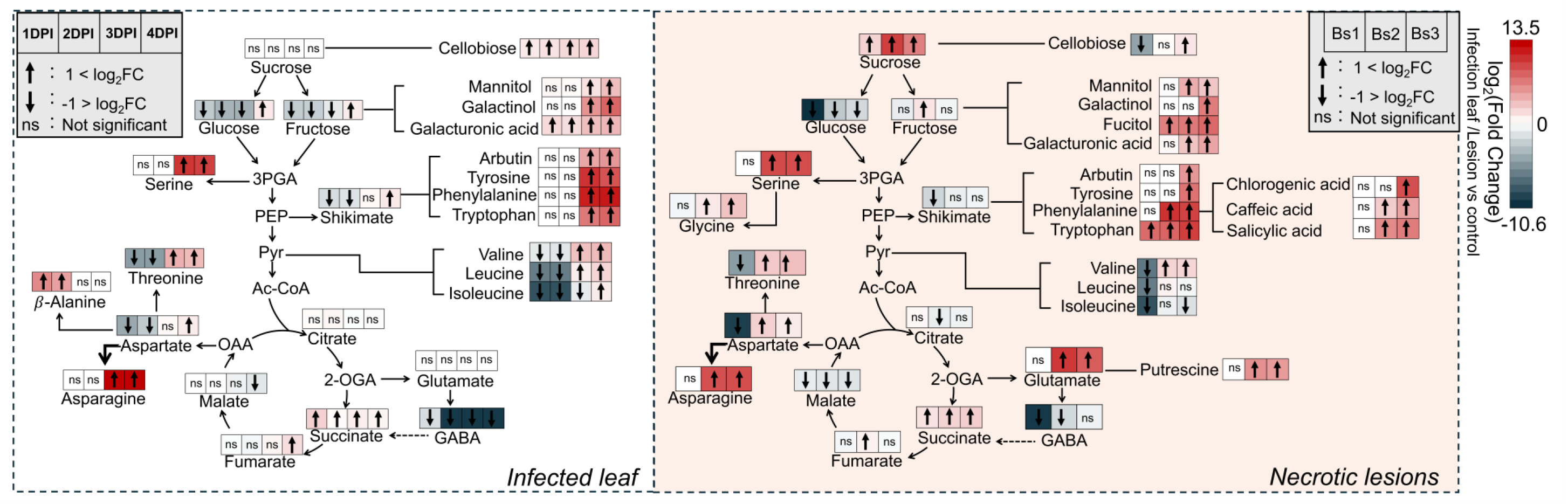
Metabolic pathway reprogramming during *Alternaria solani* infection in potato leaves. The figure summarizes changes in primary and secondary metabolites during disease progression. The left panel shows temporal changes in metabolite abundance in infected leaves at 1, 2, 3, and 4 days post-infection (DPI) relative to healthy control leaves. The right panel shows metabolite changes within the brown necrotic lesions at three stages of lesion development (Bs1, Bs2, and Bs3). Metabolite abundance is represented as log2 fold change relative to the control, with the colour scale ranging from −10.6 (dark blue, strong decrease) to +13.5 (dark red, strong increase). Within each pathway box, ↑ indicates significant upregulation (log2 FC > 1), ↓ indicates significant downregulation (log2 FC < −1), and ns indicates no significant change. Abbreviations: DPI, days post-infection; Bs, brown lesion stage.

Glucose and fructose were depleted through most of leaf infection, recovering transiently by 4 DPI; glucose was similarly depleted at Bs1-Bs2, returning to non-significance at Bs3. Sucrose was unchanged across the leaf time-course but reduced at Bs1-Bs2 in lesions. Cellobiose accumulated across all four days of leaf infection and, more modestly, at Bs3. Mannitol, galactinol, and fucitol were elevated at 3-4 DPI (leaves) and Bs1-Bs3 or Bs2-Bs3 (lesions, metabolite-dependent). Galacturonic acid rose more than 2.5-fold above control already at 1-2 DPI in leaves, and progressively from Bs2 to Bs3 in lesions. Putrescine, detected only in lesion tissue, was elevated at Bs2 and Bs3 relative to Bs1, paralleling its precursor glutamate.

### Dynamics of Glutamate-derived metabolism impacted by infection and lesions

As glutamate sits at the branch point between nitrogen assimilation, amino acid biosynthesis, and the TCA cycle, we examined the glutamate-derived metabolite pool across infection. In the plastid, the GS/GOGAT cycle assimilates NH₄⁺ into glutamine and glutamate, feeding carbon skeletons toward oxoproline and, via asparagine-synthetase-type reactions, toward asparagine and aspartate (Figure 4). In the mitochondrion, glutamate is decarboxylated to GABA, which is transaminase to succinic semialdehyde and then succinate, entering the TCA cycle via the GABA shunt. Oxoproline and asparagine followed a near-identical pattern across the leaf time-course: both were near baseline through 1-2 DPI, then rose sharply at 3-4 DPI, reaching their highest levels at 4 DPI. In lesion tissue, both were also elevated relative to control but to a smaller degree than the 3-4 DPI peak oxoproline highest at Bs1, declining through Bs2; asparagine modestly elevated across Bs1-Bs2. These increases were statistically significant relative to control (oxoproline and asparagine, p < 0.0001 at 3-4 DPI; asparagine, p < 0.01-0.0001 at lesion stages). GABA followed the opposite trajectory: highest in control, dropping sharply by 1 DPI and remaining low throughout the leaf time-course. In lesion tissue, GABA was similarly reduced at Bs1, with a partial, incomplete rise by Bs3.

**Figure 4:**
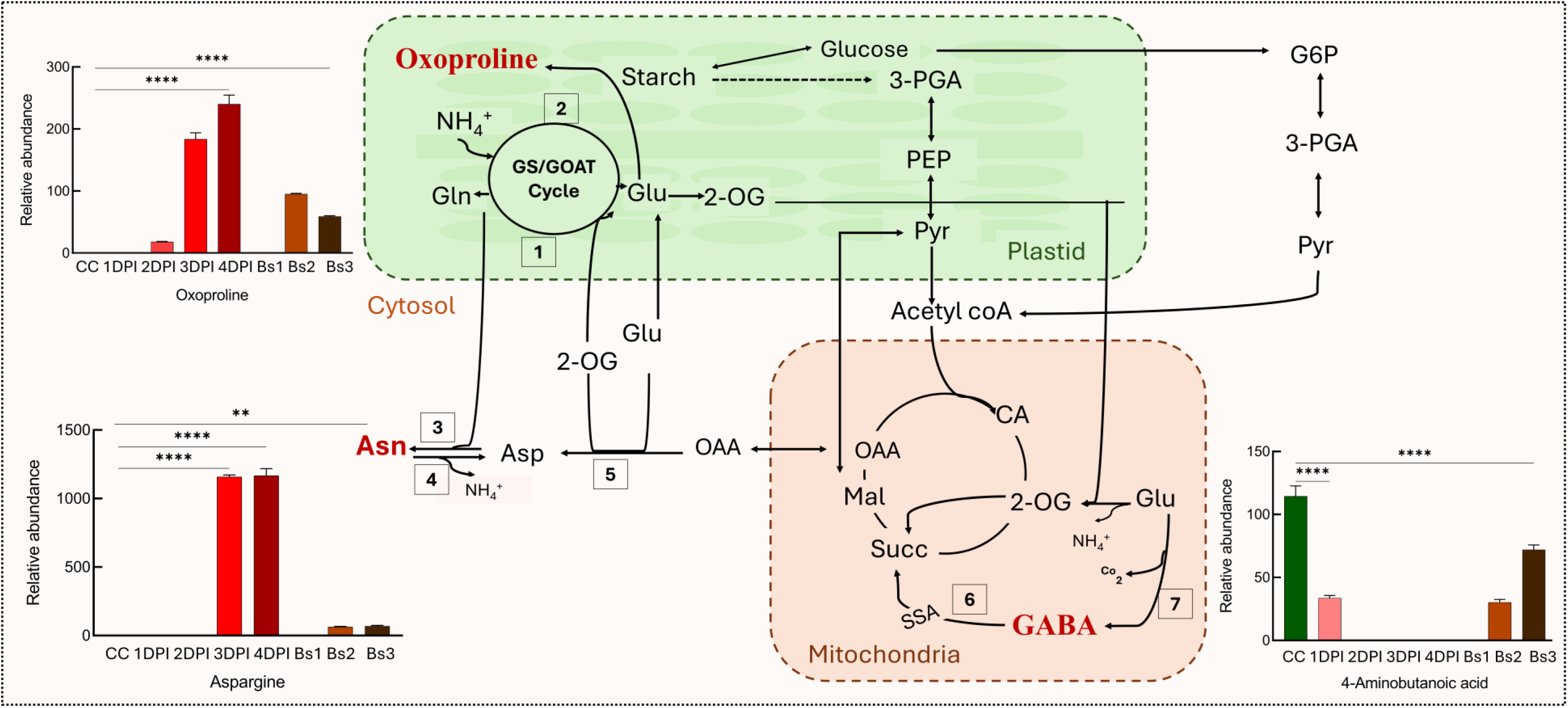
Alteration of nitrogen derived metabolites in infection and lesion contributing to defence mechanism in potato leaves under *A.solani infection*. Differential accumulation of key nitrogen-containing metabolites including glutamate, GABA, oxoproline, and asparagine was observed in infected and lesion tissues. The enzymes coded are as follows 1: ferredoxin-glutamate synthase 2: chloroplastic glutamine synthetase3: asparagine synthetase 4: asparaginase 5: cytosolic aspartate aminotransferase 6: c-aminobutyric acid transaminase 7: glutamate decarboxylase 2 [Abbreviations: CC-control leaves; DPI - Days post infection; Bs - lesion, numbers correspond to days post infection]

## 4 Discussion

### A two-axis model of infection-driven metabolic reprogramming

Taken together, the histological and metabolomic data indicate that *A. solani* infection reshapes the *Kufri Jyoti* leaf metabolome along two distinguishable axes: a gradual, cumulative shift across the whole-leaf infection time-course, and a sharper, spatially structured shift associated with lesion formation and maturation, consistent with the broader view that primary-metabolism regulation is a central, conserved component of plant-pathogen interactions (Rojas et al., 2014). The temporal sequence established in Figure 1.A symptomless control, internal lesion initiation by 1 DPI, internal lesion establishment by 2 DPI, external necrotic surge at 3-4 DPI is broadly consistent with the general infection biology described for necrotrophic *Alternaria spp*, in which localized host-cell death and internal colonisation typically precede the most dramatic external symptoms, with subsequent severity shaped by leaf age, position, and host physiological status (Brouwer et al., 2021). We note this parallel is general rather than a mechanistic demonstration specific to this cultivar. Because internal lesion formation is largely complete by 2 DPI, the sharp metabolic shifts observed at 3-4 DPI in whole-leaf profiling most plausibly reflect the consequences of an already-established infection proteolysis, nitrogen remobilisation, secondary-metabolite induction rather than the pathogen’s initial colonisation event. On this basis, 1-2 DPI can be treated as an establishment-phase window and 3-4 DPI as a consequence-phase window for interpreting the metabolomic time-course. This is corroborated by the multivariate clustering (Figure 2.A) control grouping with 1-2 DPI, followed by clear divergence by 3-4 DPI, is consistent with a delayed but cumulative metabolic response to colonisation rather than immediate wholesale reprogramming. A caveat that should be made explicit is that the lesion tissue (Bs1-Bs3) contains an increasing proportion of fungal biomass as the lesion matures, and several metabolites discussed below (sugar alcohols, organic acids) are also produced by the fungus itself or by the host-pathogen interaction.

### Suppressed TCA flux and compensatory anaplerosis

Citrate and malate were consistently depleted in both infected leaves and lesion tissues throughout disease progression (Figure 3, Supplementary Figure S.2). Similar reductions have been reported in necrotrophic infection of wild tomato (*Solanum cheesmaniae*) by *Alternaria solani*, where central-carbon-pathway metabolites were similarly perturbed (Singh et al., 2023b). In contrast, succinate accumulated specifically in lesion tissues while GABA levels declined. One possible explanation is an increased flux through the GABA shunt, which converts GABA into succinate by bypassing the 2-oxoglutarate dehydrogenase and succinyl-CoA steps of the TCA cycle. Such a metabolic adjustment could help maintain TCA cycle activity under the carbon and redox stress associated with advanced lesion development. Nevertheless, this interpretation is based only on metabolite abundance, and metabolic flux was not measured in the present study. Interestingly, fumarate accumulated at the later stages of infection in whole leaves. This increase coincided with higher tyrosine levels, raising the possibility that tyrosine catabolism contributed to fumarate production (Parker et al., 2014). Such a response may provide an alternative source of carbon to support TCA cycle metabolism as infection progresses.

### Amino acid accumulation as a defence-oriented response

The consistent accumulation of the aromatic amino acids tyrosine, phenylalanine, and tryptophan likely reflect their role as precursors for the shikimate and phenylpropanoid pathways (Figure 3, Supplementary Figure S3). This is supported by the lesion-specific accumulation of chlorogenic acid, caffeic acid, and salicylic acid, a key defense hormone, which was detected only in lesion tissue and not throughout the infected leaves. These findings suggest that the conversion of amino acid precursors into defense-related metabolites occurs primarily within the local necrotic lesion, whereas the surrounding infected leaf tissue accumulates the precursors without converting them to the same extent.

As noted above, a rising fungal-biomass fraction within maturing lesions is an alternative or additional contributor to this spatial pattern and should be considered alongside a purely host-defence interpretation. In parallel, the joint accumulation of aspartate and asparagine reflects host energy and nitrogen re-translocation strategies during stress. Aspartate accumulation aligns with its aminotransferase-mediated conversion to oxaloacetate and pyruvate, which replenishes tricarboxylic acid (TCA) cycle intermediates to sustain cellular energy production during infection (Manasseh et al., 2023). Asparagine showed a sustained increase from the earliest lesion stage onwards, suggesting a major role in nitrogen storage and redistribution during disease development. This observation agrees with its well-established function as a nitrogen transport and storage amino acid, owing to its high nitrogen-to-carbon ratio, and its accumulation is commonly associated with senescence and nitrogen remobilization in plants (Gaufichon et al., 2016). A similar pattern has been reported in other necrotrophic interactions. For example, *Botrytis cinerea* infection of susceptible tomato induces asparagine synthetase expression during host senescence, leading to asparagine accumulation that not only stores excess nitrogen in the host but may also provide a nutrient source for the pathogen (Seifi et al., 2014). In potato, asparagine and glutamine are likewise the principal amino acids involved in nitrogen transport and redistribution, particularly in actively growing and senescing tissues (Qiao et al., 2026). Together, these findings support the view that the increase in asparagine observed during *A. solani* infection reflects enhanced nitrogen remobilization as disease progresses, although the contribution of pathogen metabolism was not examined in the present study. Serine and glycine also accumulated in both infected leaves and lesion tissues. These amino acids are closely linked to photorespiration, and their increase may indicate greater photorespiratory activity as photosynthesis declines during infection (Zhang,J; et al., 2017b; Botero et al., 2018). This metabolic pattern is consistent with the chlorosis and tissue necrosis observed during lesion development, although metabolite data alone cannot confirm reduced photosynthetic activity.

### Carbohydrate remodelling and cell-wall breakdown

Glucose, fructose, and sucrose declined progressively during infection, indicating that these soluble carbohydrates were rapidly depleted to meet the increased carbon and energy demands associated with pathogen-induced stress and defence responses (Chhajed et al., 2020). Their depletion may also reflect direct uptake by *Alternaria solani*, a necrotrophic pathogen that acquires nutrients from dead and dying host tissues while secreting cell wall-degrading enzymes and other virulence factors to facilitate colonization (Brouwer et al., 2021). In contrast, cellobiose accumulated progressively throughout disease development, consistent with enhanced cellulose degradation during host cell wall disassembly, a characteristic feature of necrotrophic infection. Gluconic acid also increased markedly during lesion development. Because gluconic acid is a common product of fungal glucose oxidase-mediated oxidation of glucose (Ramachandran et al., 2006), its accumulation may partly reflect fungal metabolism. However, as mature lesions contain an increasing proportion of fungal biomass, the observed increase in gluconic acid should be interpreted as arising from combined contributions of both pathogen and host metabolism rather than representing a host-specific metabolic response. Sugar alcohols, including mannitol and galactinol, together with the lesion-specific accumulation of fucitol, became more abundant during the later stages of infection. Their delayed increase indicates that these metabolites are associated with the response to extensive tissue damage, where they may help protect cells by maintaining osmotic balance and scavenging reactive oxygen species (ROS), rather than contributing to the initial defence response. The particularly early and pronounced rise in galacturonic acid, a monomeric constituent of pectin, is consistent with pectin degradation and cell wall remodelling from the earliest stages of infection (Prade et al., 1999). Because pectin breakdown products containing galacturonic acid can act as elicitors of plant defence responses (González & Allen, 2003), this rise may represent both a symptom of cell wall damage and defence response.

### Glutamate as a metabolic hub linking nitrogen storage and defence signalling

Glutamate is a key metabolite that connects nitrogen assimilation, nitrogen storage, and the GABA shunt pathway. Changes in these pathways provide insight into how potato leaves redistribute nitrogen during disease progression. The marked increase in oxoproline and asparagine at 3-4 DPI (Figure 4) suggests that these amino acids act as important nitrogen storage compounds during the later stages of infection. As disease progresses and protein breakdown increases, the released ammonium and amino nitrogen are likely converted into oxoproline and asparagine rather than remaining as free ammonium, which can be toxic to plant cells. Oxoproline, alongside other nitrogen-rich metabolites, is additionally implicated in osmotic adjustment and cellular protection under stress (Ward et al., 2010), so its accumulation may serve a dual role as both a nitrogen sink and a stress-protective solute as tissue integrity declines late in infection. Asparagine accumulated strongly in potato during *Alternaria solani* infection, particularly at 3-4 DPI (Figure 4), suggesting enhanced nitrogen remobilization as disease progression advanced. A similar increase in asparagine has been reported in tomato leaves infected with *Pseudomonas syringae*, where its accumulation was associated with the induction of a cytosolic glutamine synthetase isoform (Pérez-García et al., 1998; Olea et al., 2004; Lea et al., 2007). However, because these studies involve different host species and pathogens, the mechanisms underlying asparagine accumulation may differ and cannot be inferred from the present data. In contrast to oxoproline and asparagine, GABA showed a different pattern during disease progression. GABA remained lower than the control throughout infection and the early lesion stages, with only a partial recovery at the late lesion stage. This decrease may have two possible explanations. First, the pathogen may utilize host-derived GABA as a source of carbon or nitrogen or suppress GABA accumulation to weaken host defence responses, as suggested for several plant pathogens (Tarkowski & Van den Ende, 2020). Second, GABA may be rapidly converted into succinic semialdehyde and subsequently succinate through the GABA shunt. This would reduce GABA levels while contributing carbon to the TCA cycle, consistent with the increased abundance of succinate-related metabolites observed in lesion tissues. Similar changes in GABA metabolism have been reported during interactions between Solanaceous plants and necrotrophic fungi, including *Botrytis cinerea*, *Rhizoctonia solani*, and *Ralstonia solanacearum* (Seifi et al., 2013; Tarkowski & Van den Ende, 2020; Rani et al., 2021; Liu et al., 2024; Zarbakhsh et al., 2025). Together, these findings suggest that GABA metabolism is an important component of the plant response to necrotrophic infection, although the underlying mechanisms were not directly investigated.

## 5. Conclusion

Comprehensive metabolite profiling revealed that *Alternaria solani* infection causes extensive metabolic changes in potato leaves, affecting pathways involved in carbohydrate metabolism, energy production, nitrogen remobilization, and secondary metabolism. The progressive changes observed in glycolysis, the TCA cycle, β-oxidation, GS/GOGAT, and the shikimate pathway indicate that infection is accompanied by substantial metabolic reorganization as the plant responds to increasing disease severity. In contrast, the early activation of inositol phosphate and glutathione metabolism, together with the later accumulation of amino acids such as aspartic acid, threonine, and phenylalanine, suggests the activation of defence-related processes during disease progression. The lesion-specific analysis showed that necrotic lesions are metabolically distinct from the surrounding infected leaf tissue. At the earliest lesion stage (Bs1), reduced levels of oxoproline and asparagine point to disruption of nitrogen metabolism, while the low abundance of GABA suggests altered carbon and nitrogen partitioning. As lesions developed (Bs2-Bs3), several defence-associated metabolites accumulated, indicating continued metabolic adjustment within infected tissues. These differences between whole infected leaves and lesion tissues demonstrate that disease progression is accompanied by spatial as well as temporal changes in metabolism. Several metabolites, including asparagine, oxoproline, GABA, aspartic acid, and phenylalanine, emerged as potential metabolic markers associated with early blight progression. Although the present study was based on steady-state metabolite profiling, future work using stable isotope (^13^C) tracer experiments could determine how carbon is redistributed through the TCA cycle, shikimate pathway, and phenylpropanoid metabolism during infection. Moreover, the identification of key discriminatory metabolites opens new avenues for developing diagnostic markers and protective compound formulations against early blight. Integrating these findings with genome editing and conventional breeding holds promise for enhancing resistance traits and improving crop resilience against biotic stresses in sustainable agriculture.

## Acknowledgment

PDS acknowledges the SaIAFarm project and the Ministry of Education for PhD fellowship. RN acknowledges the Ministry of Education for MTech (by Research) fellowship.

## Declarations

### Ethical approval

The study complied with the ethical standards.

### Data availability

The datasets generated during and/or analysed during the current study are available from the corresponding author on reasonable request

