## Supplemental figures and tables for "*Alternaria solani* infection reprograms potato leaf metabolism and highlights potential defence and metabolic markers"

**Supplementary Table 1: The metabolite profiles of potato leaves were analysed under *A.solani* infection in three conditions: control, infected, and brown spots.**

| S.No | Metabolite name | CC | DPI1 | DPI2 | DPI3 | DPI4 | Bs1 | Bs2 | Bs3 |
|---|---|---|---|---|---|---|---|---|---|
| 1 | Valine | 12.3 | 5.9 | 5.2 | 120.2 | 159.3 | 0 | 60.1 | 56.4 |
| 2 | Leucine | 10 | 0 | 0 | 19.4 | 48.2 | 0 | 16.4 | 7.6 |
| 3 | Isoleucine | 46.7 | 0 | 0 | 23.5 | 99.7 | 0 | 28.1 | 16 |
| 4 | Serine | 0 | 0 | 0 | 161.3 | 114.4 | 0 | 77.5 | 69.8 |
| 5 | Threonine | 3.2 | 0 | 0 | 60.4 | 39.1 | 0 | 24.5 | 33.7 |
| 6 | Glycine | 19.1 | 18.4 | 14.8 | 13.5 | 18.8 | 13.6 | 46 | 176.3 |
| 7 | Aspartic acid | 93.7 | 4.4 | 14 | 124.9 | 190.9 | 0 | 525.1 | 199.8 |
| 8 | Asparagine | 0 | 0 | 0 | 1158.4 | 1166.8 | 0 | 62.8 | 68.3 |
| 9 | Glutamic acid | 0 | 0 | 0 | 0.1 | 0.1 | 0 | 150.2 | 40.7 |
| 10 | Phenylalanine | 0 | 0 | 0 | 393.6 | 483.5 | 0 | 190.3 | 102 |
| 11 | Tyrosine | 0 | 0 | 0 | 193.4 | 57.1 | 0 | 0 | 17.5 |
| 12 | β-Alanine | 0 | 7.9 | 3.2 | 0 | 0 | 0 | 0 | 0 |
| 13 | Tryptophan | 0 | 0 | 0 | 14.6 | 29.4 | 24.1 | 30.2 | 122.6 |
| 14 | Hydroxylamine | 28.7 | 23.9 | 24.9 | 14.9 | 21.1 | 28.6 | 15.8 | 18.4 |
| 15 | Oxoproline | 0 | 0 | 18.1 | 183.6 | 240.2 | 0 | 95.1 | 59.1 |
| 16 | Aminobutanoic acid | 114.4 | 33.7 | 0 | 0 | 0 | 0 | 30.3 | 72 |
| 17 | Trihydroxybutyric acid | 14.6 | 118.6 | 76 | 18.9 | 12.5 | 73.4 | 18.6 | 32.6 |
| 18 | Urea | 96.2 | 126.2 | 127 | 62.4 | 55.2 | 137.8 | 56.4 | 74.2 |
| 19 | Putrescine | 0 | 0 | 0 | 0 | 0 | 0 | 5.8 | 3.9 |
| 20 | Succinic acid | 9.4 | 51.2 | 11.6 | 17.9 | 14.8 | 27.5 | 48.3 | 58.8 |
| 21 | Fumaric acid | 20.4 | 18.5 | 15 | 31.9 | 68.9 | 15.1 | 35.2 | 17.3 |
| 22 | Malic acid | 318 | 311.9 | 340.2 | 190.7 | 136 | 160.4 | 130 | 121.9 |
| 23 | Citric acid | 215.8 | 257.2 | 308.8 | 201.3 | 193.9 | 297.3 | 119.9 | 140.8 |
| 24 | Ethanolamine | 61 | 27.4 | 21 | 61.2 | 47.2 | 44.7 | 47.9 | 89.7 |
| 25 | Ribose | 0 | 0 | 0 | 0 | 0 | 2.4 | 106.2 | 108.3 |
| 26 | Arabinose | 34.7 | 46.1 | 46.5 | 64.2 | 55.9 | 34 | 124.4 | 186 |
| 27 | 1,5-Anhydroglucitol | 61 | 0 | 0 | 4 | 8.9 | 0 | 0 | 0 |
| 28 | Fucitol | 0 | 0 | 0 | 0 | 0 | 11 | 13.8 | 26.4 |
| 29 | Gluconic acid | 0 | 44.1 | 33.7 | 45.1 | 68.6 | 42 | 323.8 | 188.9 |
| 30 | Galactaric acid | 0 | 0 | 0 | 0 | 0 | 3.1 | 12 | 15.7 |
| 31 | Fructose | 134.5 | 22 | 23.7 | 61.5 | 271.5 | 73.1 | 378 | 98.5 |
| 32 | Galactose | 7.9 | 14.3 | 19.4 | 6.5 | 31.7 | 1.7 | 305.1 | 420.6 |
| 33 | Glucose | 158 | 24.4 | 11 | 9.9 | 357.7 | 0 | 23.6 | 42 |

| | | | | | | | | | |
|---|---|---|---|---|---|---|---|---|---|
| 34 | Mannitol | 8.4 | 11.5 | 9.9 | 24.4 | 92.3 | 7.5 | 174.3 | 69.4 |
| 35 | Galactinol | 0 | 0 | 0 | 5.3 | 29.1 | 0 | 0 | 12.5 |
| 36 | Myo-Inositol | 63.2 | 12.4 | 11.7 | 32.1 | 125 | 9.8 | 103.5 | 71.9 |
| 37 | Dulcitol | 20.8 | 0 | 0 | 0 | 0 | 8.8 | 5.8 | 63.4 |
| 38 | Galacturonic acid | 3.3 | 12.4 | 11.2 | 36.1 | 46.3 | 5.3 | 33.3 | 101.1 |
| 39 | Cellobiose | 2.7 | 6.1 | 7.8 | 10.1 | 11.8 | 0.2 | 3.1 | 9.8 |
| 40 | Maltose | 0 | 26.4 | 21.2 | 10.5 | 16.3 | 7.4 | 20.1 | 146.7 |
| 41 | Mannobiose | 0 | 73.9 | 23 | 18.2 | 7.3 | 0.1 | 16.3 | 55.1 |
| 42 | Sucrose | 0 | 0 | 0 | 0 | 0 | 0.5 | 116.9 | 11.4 |
| 43 | Glycerol monostearate | 66.5 | 35.2 | 32.3 | 66.7 | 92.3 | 16.9 | 54.7 | 81.9 |
| 44 | Shikimic acid | 17.8 | 5.2 | 6.8 | 23.9 | 40.5 | 4.3 | 9.8 | 11.8 |
| 45 | Salicylic acid | 0 | 0 | 0 | 0 | 0 | 0 | 7.4 | 11.9 |
| 46 | Caffeic acid | 0 | 0 | 0 | 0 | 0 | 0 | 0.7 | 3.1 |
| 47 | 5-O-Coumaroyl-D-quinic acid | 0 | 0 | 0 | 0 | 0 | 0 | 0 | 6.8 |
| 48 | Arbutin | 0 | 0 | 0 | 2.6 | 4.9 | 0 | 0 | 4.4 |
| 49 | Chlorogenic acid | 0 | 0 | 0 | 0 | 0 | 0 | 0 | 63 |
| 50 | Palmitic Acid | 231.4 | 173.1 | 169.9 | 182.1 | 191 | 131.7 | 119.5 | 140.6 |
| 51 | Stearic acid | 102.4 | 66.2 | 66.7 | 87.7 | 103 | 45 | 61.5 | 80.6 |
| 52 | Monopalmitin | 106.8 | 63.5 | 62.6 | 78.2 | 91.8 | 47.9 | 67.8 | 89.7 |
| 53 | Aucubin | 0 | 0 | 0 | 0 | 0 | 0 | 7.3 | 42.1 |
| 54 | Quininic acid | 66.4 | 42.1 | 38.3 | 74.3 | 99.2 | 25.8 | 35.3 | 88 |
| 55 | Propanedioic acid | 30.9 | 43.9 | 43.7 | 29.6 | 37.6 | 22.6 | 47.3 | 62.7 |
| 56 | Glyceric acid | 0 | 78.8 | 67.4 | 81.3 | 60.2 | 0 | 66.4 | 43.8 |
| 58 | Oxalic acid | 47.7 | 16 | 48.6 | 160.1 | 157.8 | 0 | 5.9 | 15.8 |

Values are normalised by internal standard and mean of (n=4). Abbreviations: CC-control leaves; DPI: Days post infection; bs: Lesion developed; numbers correspond to days post infection.

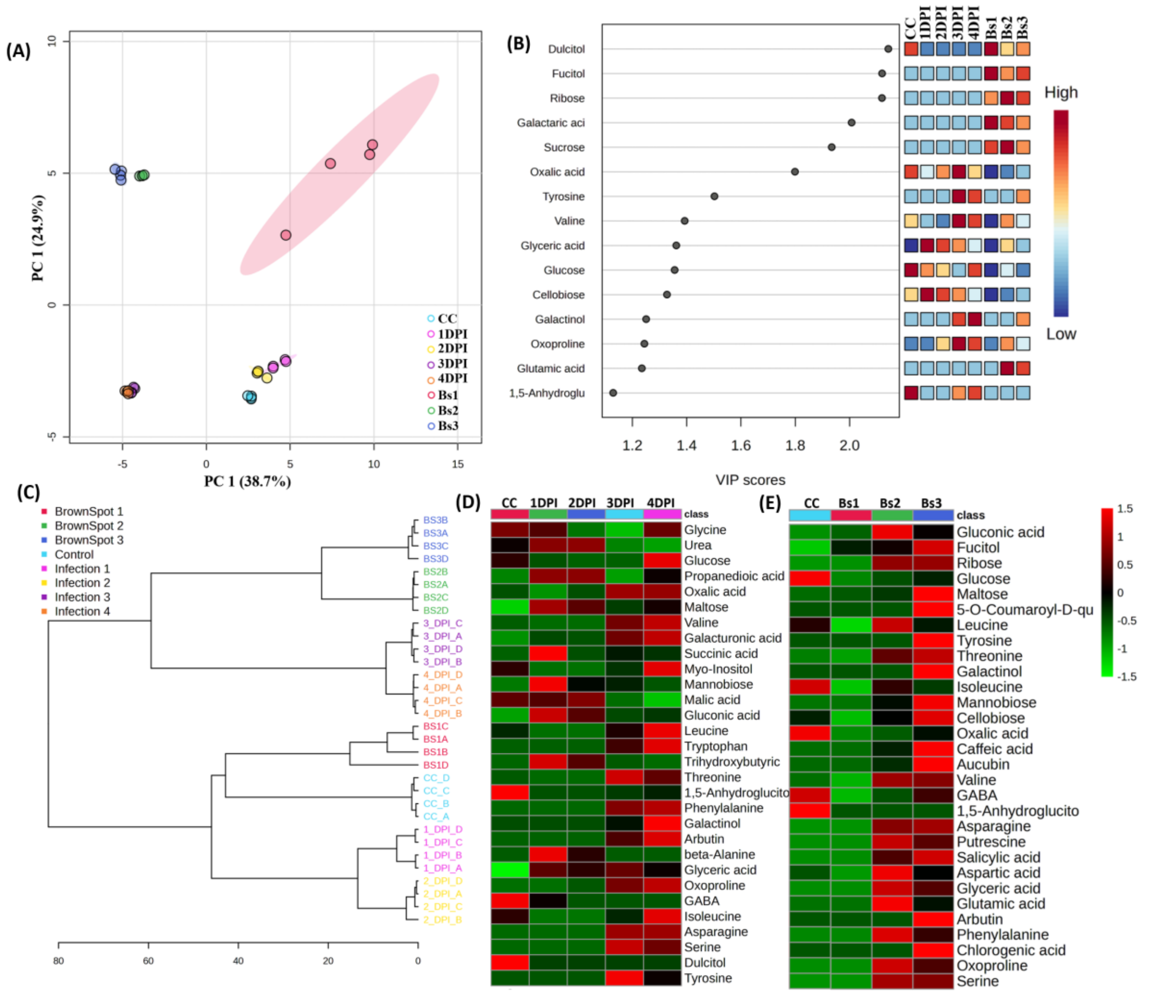


**Supplementary Figure S 1: Comparative multivariate analysis in potato leaves under *A.solani* infection. A)** PCA analysis also showed clear separation among the treatments on the variability across the two principal components (PCs) 1 and 2, respectively. **B)** The VIP score plots display the significance of 20 identified metabolites with a VIP score greater than 1.A trichromatic scale consisting of blue, yellow, and red colors is employed to visually represent the extent of a metabolite's contribution to metabolic variation in different treatment conditions **C)** Dendrogram reveal that control, lesions at day 1 and infection at day 1 and 2 after infection modulate their metabolome likewise under *A.solani* infection, whereas infection at day 3 and 4 and lesion at day 2 and 3 modulate similar pattern. **D)** Heatmap showing the metabolite difference in control and progression of infection. **E)** Heat maps showing the metabolite differences in control and collected lesions, levels are presented as green to red scale Abbreviations: CC-control leaves; DPI: Days post infection; Bs: Lesion developed; numbers correspond to days post infection

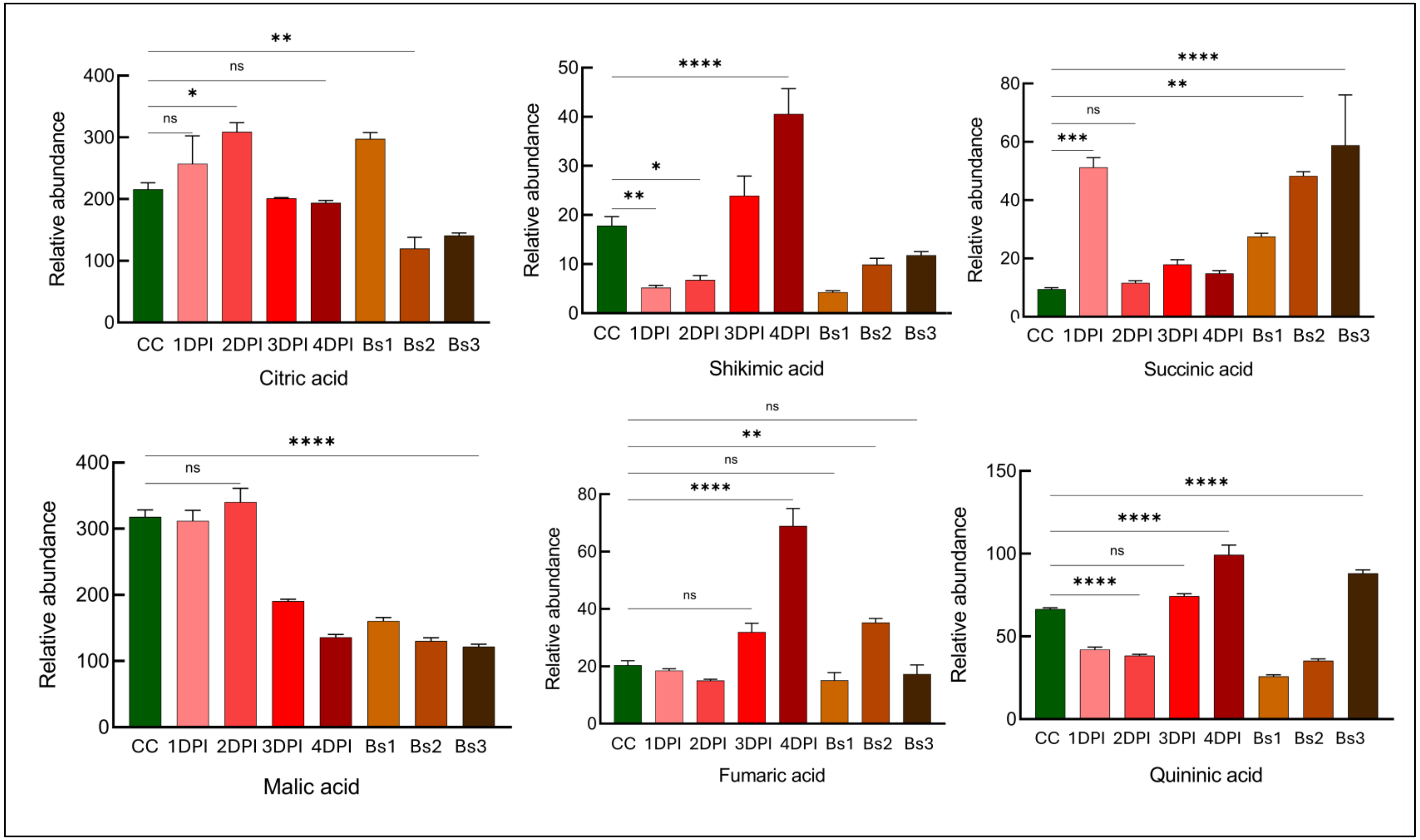


**Supplementary Figure S2:** Relative quantitative metabolic variations in central metabolites in potato leaves under *A.solani* infection in 3 condition i.e. control, infected and lesion. Abbreviations: CC-control leaves; DPI: Days post infection; bs: Lesion developed; numbers correspond to days post infection

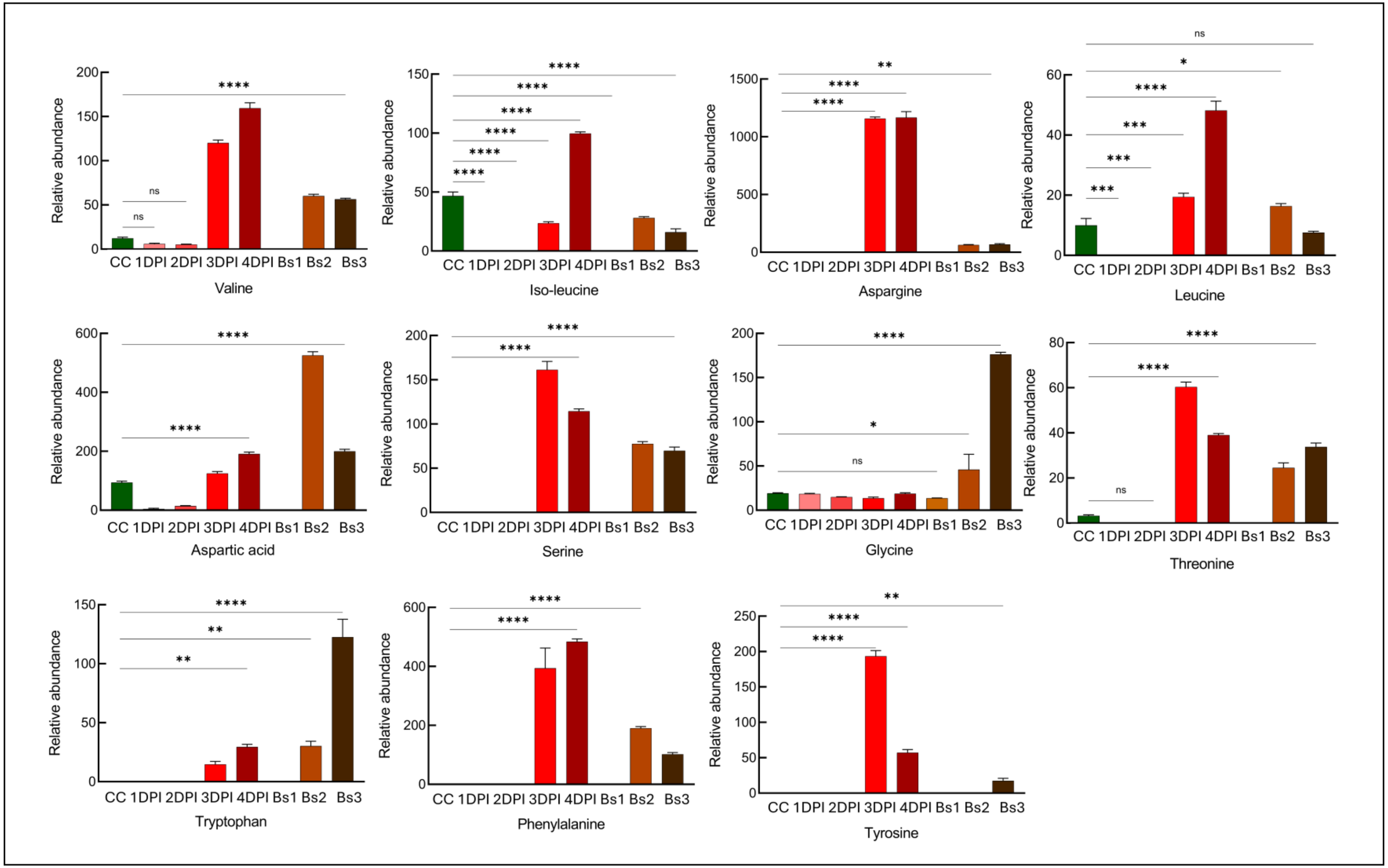


**Supplementary Figure S3:** Relative quantitative metabolic variations in amino acids in potato leaves under *A.solani* infection in 3 condition i.e. control, infected and lesions. Abbreviations: CC-control leaves; DPI: Days post infection; bs: Lesion developed; numbers correspond to days post infection
